# Simulating semantic dementia in a brain-constrained model of action and object words learning

**DOI:** 10.1101/2022.03.03.482066

**Authors:** Aleksei Efremov, Anastasia Kuptsova, Thomas Wennekers, Yury Shtyrov, Boris Gutkin, Max Garagnani

## Abstract

The nature of semantic knowledge – conceptual information stored in the brain – is highly debated in the field of cognitive science. Experimental and clinical data specify various cortical regions involved in the processing of meaning. Those include semantic hubs that take part in semantic processing in general as well as sensorimotor areas that process specific conceptual categories according to their modality. Biologically inspired neurocomputational models can help adjudicate between different theories about the exact roles of those regions in the functioning of the semantic system.

Here, we used an existing neuroanatomically constrained model of frontotemporal brain areas implicated in language acquisition and grounding. We adapted it to replicate and explain the effects of semantic dementia on word processing abilities. Semantic dementia is a disease characterized by semantic knowledge deterioration that correlates with neural damage in the anterior temporal lobe. The behavior of our model is in accordance with clinical data – namely, word recognition performance decreases as SD lesions progress, whereas word repetition abilities remain preserved, or are less affected. Furthermore, our model makes novel predictions about category-specific effects of SD – namely, our simulation results indicate that word processing should be more impaired for object-than for action-related words, and that white matter degradation should lead to more severe consequences than the same proportion of grey matter degradation.

The present results provide a mechanistic, cortical-level explanatory account of a range of language impairments as observed in clinical populations during the onset and progress of semantic dementia.

## Introduction

Semantic knowledge is information about the meaning of concepts and words accumulated by an individual throughout their lifetime (1, 2). We constantly use it to behave properly in different situations, express our thoughts, understand others, and for many more purposes. Difficulties experienced by people with semantic memory impairments can be of a diverse nature — they can fail to recognize or produce speech (3, 4); they can forget how to use ordinary things, for example, an umbrella (5); or they can fail to understand and explain the source of the pain that they feel (4). This shows the importance of understanding the nature of such impairments and the nature of semantic knowledge itself.

Although conceptual information processing in the brain has been one of the central research topics in recent decades, its neural substrate, processing mechanisms, or even the substance of conceptual information remain an open debate (6). One line of reasoning posits that the sensorimotor system plays only a secondary role in semantic representations (7, 8) and that it is important to have one higher-order brain area where all concepts are stored in an amodal way (9, 10). Other groups claim that when we process a concept, our brain simulates the perceptual experience of the interaction with this concept (11). Finally, multiple scholars propose hybrid theories that suggest both the necessity of symbol grounding in perceptual and motor experiences and the existence of ‘higher order’ convergence areas that link together experiences from different modalities (1, 12–14).

A huge amount of evidence on the functioning of the semantic system comes from clinical studies. Semantic dementia (SD) is an exemplar semantic disorder that gave birth to numerous insights into conceptual information processing mechanisms (5, 15, 16). It is characterized by deterioration of semantic knowledge, the degree of which correlates with the severity of anterior temporal lobe (ATL) atrophy (3, 4, 17). Furthermore, according to some computational models of SD (15, 18) and transcranial magnetic stimulation (TMS) studies (19), ATL atrophy is considered as the cause of impairments of semantic memory. Several researchers have suggested that this evidence supports the existence of an amodal core of concept representations and diminishes the importance of concepts embodiment (9, 20, 21). This line of thinking stems from the observation that semantic memory decline in SD patients is not concept selective (i.e. different semantic categories are affected) and ATL is not part of the sensory and motor systems, therefore ATL may be the core place where concepts from different categories are stored in an amodal way.

On the other hand, amodal representations cannot fully explain the data on the correlation between activation of different cortical sites and comprehension of specific conceptual categories. Patients with lesions in the motor and premotor cortex often suffer from impaired comprehension of action-related words compared with object-related ones, while patients with lesions in the visual cortex have problems with comprehension of object-related words (for a review, see (1, 22). Furthermore, it was shown that TMS of the hand motor areas could enhance comprehension of the hand-related words, while stimulation of the leg areas could have a similar effect on the leg-related words (23). Other TMS studies show that comprehension and retrieval of different categories of concepts (e.g., animals versus tools) correlate with different cortical stimulation sites (in this example, occipital and inferior temporal areas versus premotor areas, accordingly; for reviews see (24, 25). This line of reasoning posits that the neuronal networks integrating sensory and motor patterns that emerge as a result of concomitant action and perception experiences during word meaning and concept acquisition could be the basis for semantic representations. More specifically, Kiefer and Pulvermüller (1) suggest that concept representations are distributed over modality-specific sensorimotor areas and non-specific (or modality general) “connector hubs”, which provide the neuroanatomical “bridge” between the different modality-specific cortices. Brain connectivity data show that relevant primary motor and sensory areas are not directly linked, but can only communicate by virtue of mediating areas (connector hub). Therefore, conceptual representations must necessarily be distributed over modality-specific sensorimotor areas and general connector hubs. They argue that this hypothesis follows directly from the perceptual origin of semantic knowledge acquisition and the neurobiological properties of the brain (14, 25). When we acquire the meaning of a new concrete word or concept, this usually occurs through different perceptual modalities simultaneously (for example, a mother teaches her child ‘This is an apple’ by showing an apple, so in this case, auditory and visual perception occurs at the same time). This co-occurrence could establish robust links between neural networks subserving different perceptual and motor modalities through Hebbian learning mechanisms (26).

In this study, we extend a neuroanatomically constrained computational model of word acquisition and grounding (14) to reflect neural mechanisms associated with semantic dementia. Using this model, we attempt to explain existing data about SD syndrome, namely, why word recognition abilities decline with an increase in the disease severity while word repetition abilities remain relatively preserved. In the extended model, we implement two types of neural anomalies reported to be associated with SD —damage to the grey matter and white matter of the ATL (3, 4, 17) — and compare the model dynamics resulting from these two alterations. We also compare the degradation dynamics of object-related words with the degradation dynamics of action-related words during the simulated SD progression.

## Methods

### Overview of the model

To address the question of ATL involvement in the process of semantic knowledge deterioration during SD, we implemented a neurobiologically constrained model that replicates selected cortical areas and their connectivity. The model simulates 12 areas of the left hemisphere cortex (Fig.1). Six of them are perisylvian areas that are involved in language production and comprehension. The other six areas are believed to be involved in transferring and processing semantically relevant information during word meaning acquisition and comprehension / recognition – in the remainder of this article, they are referred to as extrasylvian areas (9, 11, 27, 28). The model is further divided into four modality-specific ‘zones’, with each zone containing three cortical areas: a primary sensorimotor cortex and adjacent ‘higher’ secondary and multimodal regions that have strong neuroanatomical links with this primary cortex (see subsection ‘Connectivity of the Simulated Brain Areas’ below).

**Fig. 1.**
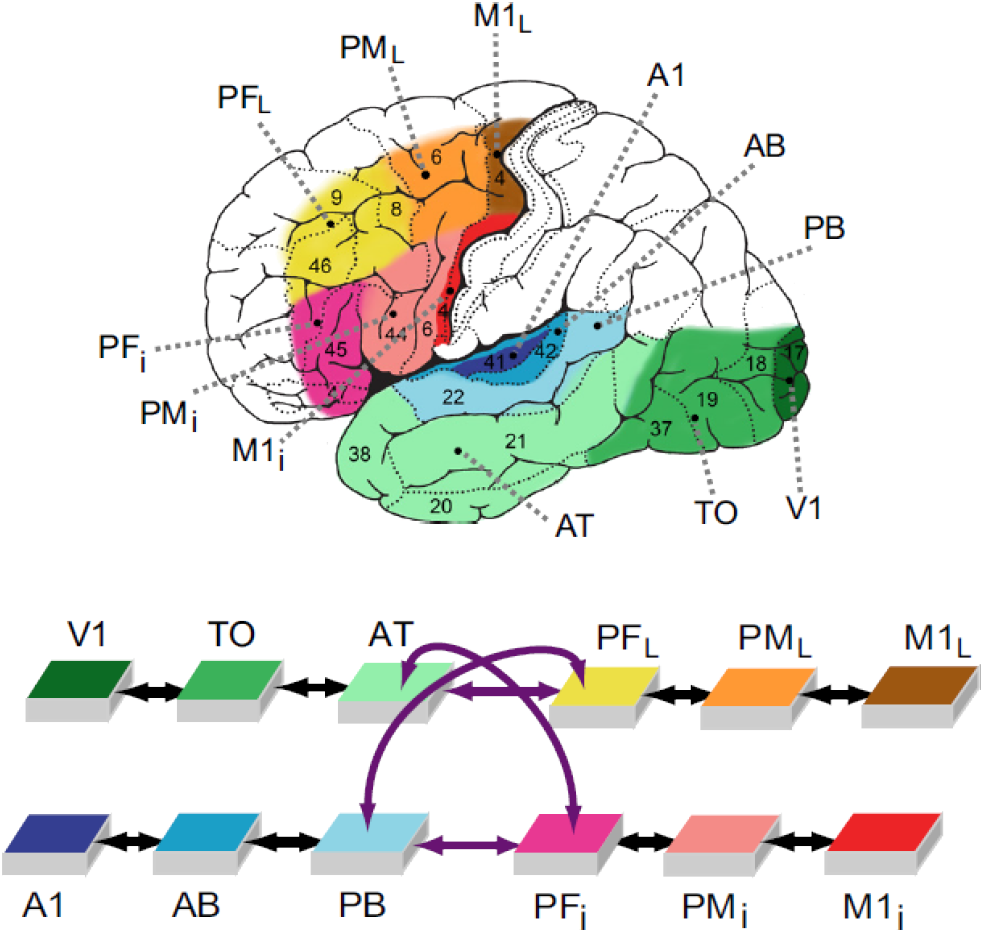
Macrostructure of the model. Depicted here are 12 areas of the network and corresponding brain areas (coded by color). There are four ‘zones’ of three different modalities, with three areas in each zone. The ‘auditory’ zone comprises the superior and lateral auditory areas: primary auditory area (A1), auditory belt (AB) and auditory parabelt (PB). The ‘visual’ zone comprises the inferior temporo-occipital areas: primary visual area (V1), temporo-occipital area (TO) and anterior temporal area (AT). The motor cortex is represented by two zones – one corresponds to the articulatory movements, and the other (which we refer to as just the ‘motor’ zone) to nonarticulatory movements. The ‘articulatory’ zone comprises the inferior frontal areas: inferior primary motor area (M1i), inferior premotor area (PMi) and inferior prefrontal area (PFi). The motor zone comprises the superior-lateral frontal areas: lateral primary motor area (M1L), lateral premotor area (PML), and dorsolateral prefrontal cortex (PFL). We differentiate between the perisylvian cortex (auditory and articulatory areas) and the extrasylvian cortex (visual and motor areas). Black arrows indicate connections between adjacent areas, while the purple arrows indicate long-distance connections.

Each area consists of two neuronal layers — an excitatory (e-cells) layer and an inhibitory (i-cells) one — with 625 (25×25) cells in each layer. Each i-cell corresponds to exactly one e-cell; a combination of an e-cell and an i-cell reflects approximately one cortical column that consists of pyramidal excitatory neurons and inhibitory interneurons. Cells are modeled as graded-response neurons (see Appendix (Detailed Model Implementation) for specifications).

Functional and structural features of the model reflect well-documented properties of the human cortex such as:

1. known structure of the neuroanatomical links between the modeled brain systems;
2. sparse, patchy, and topographic between-and within-area connections, with the probability of the existence of a synaptic link between two cells falling off with their distance (29, 30) (see Appendix (Detailed Model Implementation) for specifications);
3. local lateral inhibition (Fig.2) and area-specific global regulation mechanisms (31);
4. Hebbian learning mechanisms that represent phenomena of long-term potentiation and depression (32);
5. neurophysiological dynamics of single cells including temporal summation of inputs, sigmoid transformation of membrane potentials into neuronal outputs, and adaptation (33);
6. presence of uniform white noise (simulating spontaneous baseline neuronal firing) in all parts of the network at all time points (34).

**Fig. 2.**
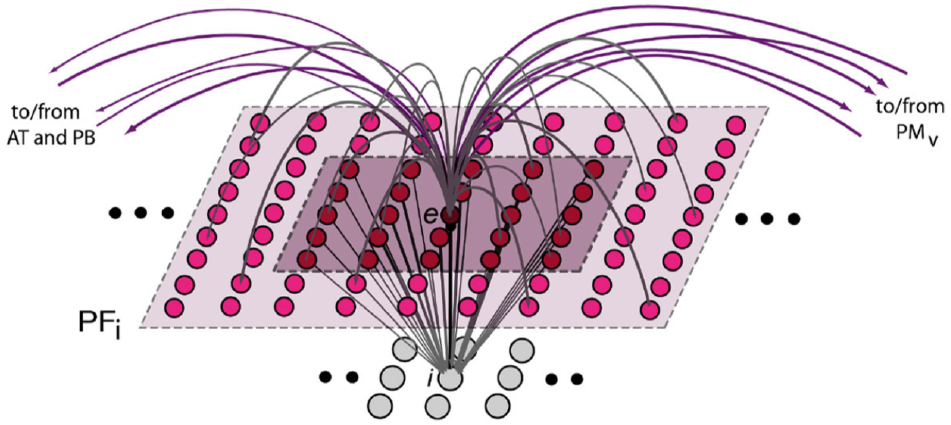
Schematics of microconnectivity of one of the 7,500 excitatory neural elements (labelled ‘e’). Within-area excitatory links (in grey) to and from ‘cell’ e are random and sparse, and limited to a local (19×19) neighborhood (light-pink shaded area).Lateral inhibition between e and neighboring excitatory elements is realized as follows: the underlying cell ‘i’ inhibits e, while its activity depends on the total excitatory input it receives from the 5×5 neighborhood around e (dark-purple shaded area); by means of analogous connections (not depicted), e can inhibit its neighbors.

### Connectivity of the simulated brain areas

Neuroanatomical studies show that adjacent cortical areas tend to be connected with each other (35, 36). We implemented such connections in all of the four zones of our model, between (1) inferior frontal areas PFi–PMi–M1i; (2) dorsolateral frontal areas PFL–PML–M1L (see also (37–40); (3) superior and lateral auditory areas A1–AB–PB (41–43); and (4) inferior temporo-occipital areas V1–TO–AT (44, 45).

Implementation of long-distance cortico-cortical links (purple arrows in Fig.1) that connect distant areas with each other is abundantly supported by evidence. Arcuate and uncinate fascicles provide the connections between anterior, inferior, and posterior-superior parts of the temporal cortex (areas AT and PB) and the inferior prefrontal cortex (PFi; (46–55). The extreme capsule connects the dorsolateral prefrontal cortex (PFL) to anterior and inferior temporal regions (AT; (56–58) and to the superior temporal cortex (PB; (54, 56, 59–61).

### Learning words of different semantic categories

As in (14), the model was taught to differentiate between two semantic categories: action-and object-related words. The model simulates the acquisition of a word having object-related semantics by means of co-experiencing and associating auditory, articulatory, and visual patterns, as inputs to A1, M1i, and V1 are presented simultaneously during the learning phase (Fig.3A). To teach action-related semantics, input patterns are provided to auditory (A1), articulatory (M1i), and motor (M1L) primary areas (Fig.3B). Presenting a “pattern” as input involves activating 19 pre-specified cells in each of the aforementioned areas. The set of cells to be activated was randomly assigned for each word. Six word patterns for each of the two semantic categories were generated and taught; therefore, the model learned 12 different word patterns.

**Fig. 3.**
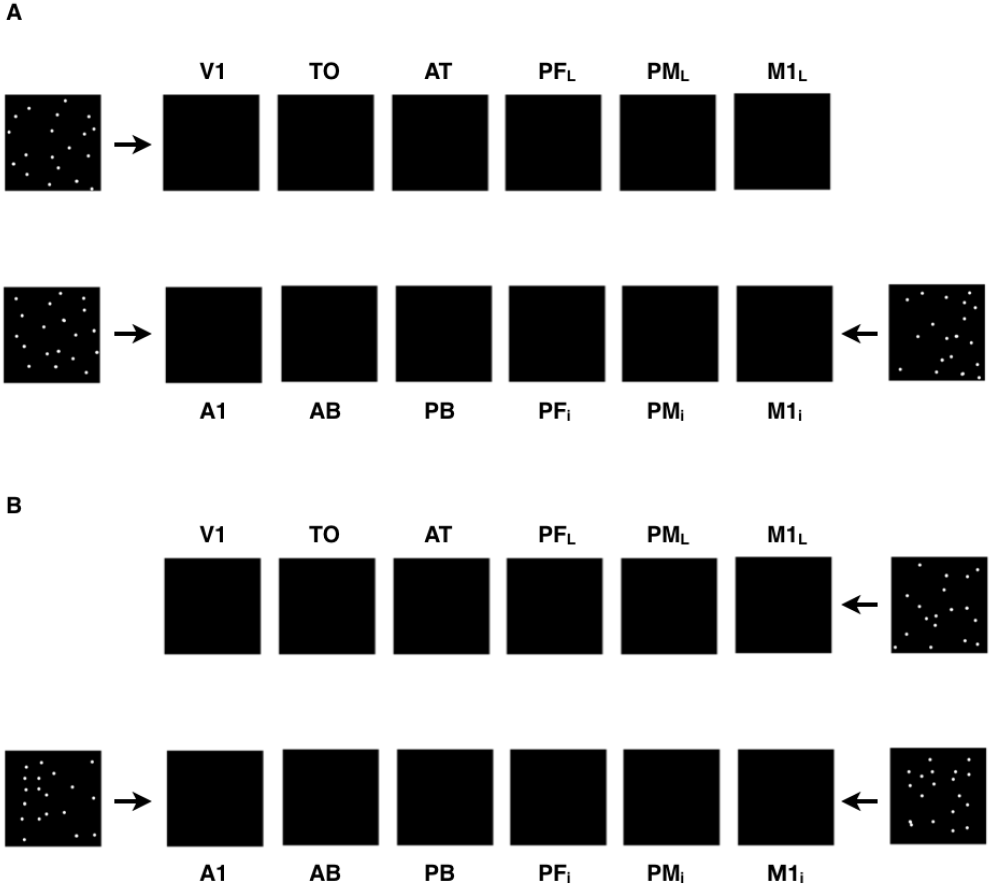
Example of activity patterns presented to the network’s primary areas to simulate word learning. A – The acquisition of object-related words was simulated by providing input patterns simultaneously to the model correlates of the auditory (A1), articulatory (M1i) and visual (V1) areas; B – Similarly, the simulation of action-related words acquisition involved presenting concomitant inputs to the auditory (A1), articulatory (M1i) and motor (M1L) areas. See text for details.

During the learning procedure, each word pattern is presented for 3000 trials. One trial lasts 16 time-steps. The next trial starts as soon as the activity in the network falls below the threshold but not earlier than 30 time-steps from the previous trial end. Trials are randomly shuffled. To reflect larger variability of activity in noninvolved primary areas (V1 for action-related words and M1L for object-related words), a randomly generated 19-cell noise pattern is provided to them in each trial. Finally, in addition to the uniform white noise constantly present in all areas of the network (simulating spontaneous neuronal activity), four primary areas are presented with additional “environmental” white noise both during training and testing.

### Implementation of semantic dementia

Neuronal, anatomical and functional changes during SD can be described as a bilateral and asymmetric pattern of progressive atrophy and hypometabolism in white and grey matter (3, 4, 17). Grey matter changes are detected in the anterior temporal lobes — superior, middle, inferior temporal gyri, fusiform gyrus, temporal pole, parahippocampal gyrus — and progress to the basal ganglia and the medial orbitofrontal cortex during the course of the disease (for a review, see (17, 62). Changes in white matter are detected in regions that are connected to or are adjacent to the temporal lobes: left inferior fronto-occipital fasciculus; uncinate fasciculus and inferior longitudinal fasciculus bilaterally (for a review, see (17, 62). The most severe and the most robust atrophy in SD patients is detected in the anterior temporal pole and ventral parts of the ATL (20).

Following that evidence, we apply two types of degradation to simulate SD — grey matter degradation (GM SD) and white matter degradation (WM SD). To simulate GM SD we inactivate e-cells in the AT area (Fig.4A) and to simulate WM damage we remove connections to and from those cells (both within-area and between-areas; Fig.4B). We also use three severity levels for each degradation type — 30%, 60%, and 90% loss of matter — to simulate effects of the progressive nature of this disorder (3, 4, 17).

**Fig. 4.**
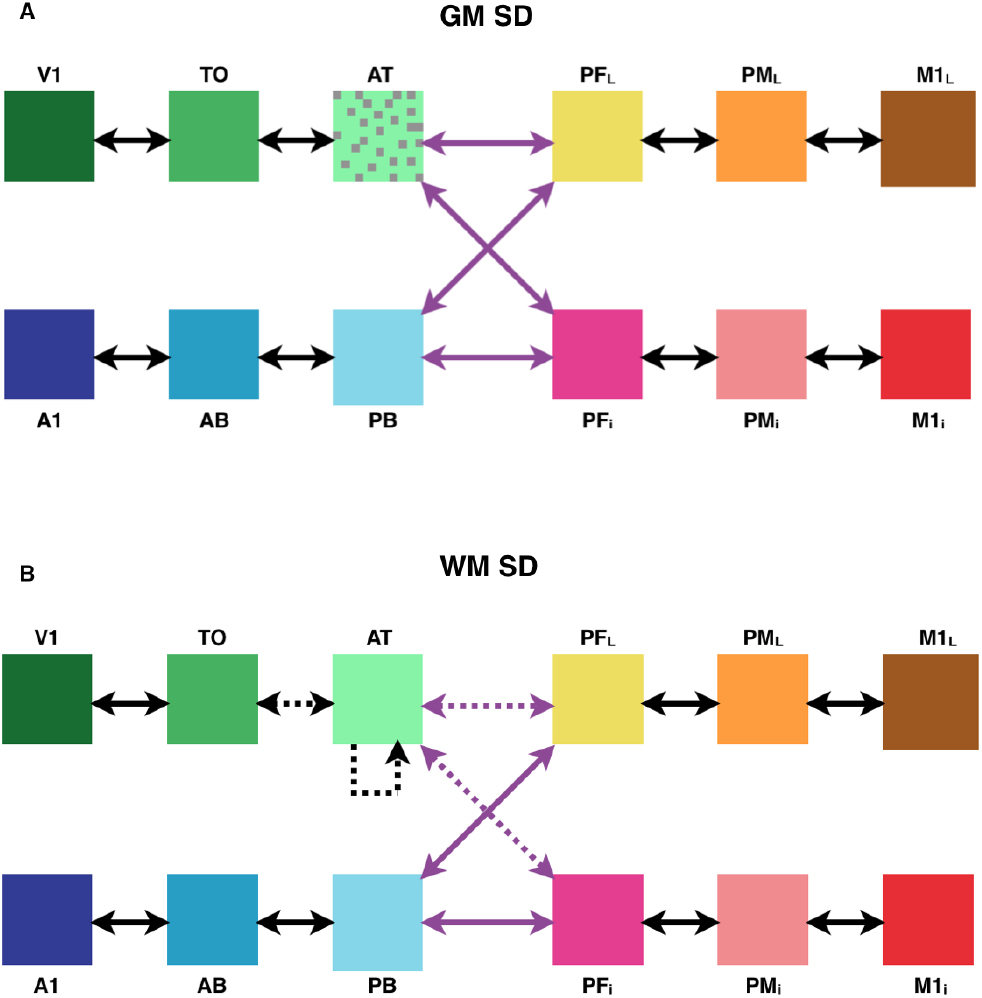
Simulated semantic dementia lesions. Schematic representation indicating the type and location of the model lesions applied to simulate grey matter semantic dementia (GM SD, A) and white matter semantic dementia (WM SD, B). Dotted area represents inactivation of some portion of e-cells in AT area. Dashed lines represent inactivation of some portion of corresponding links.

First, we trained 13 nets using the learning procedure described above, then we copied the resulting trained nets, and then the copies underwent one of the two degradation types (GM SD or WM SD) of the particular severity level (30%, 60%, or 90% loss of matter). Thus, we replicated the situation with SD patients who acquired semantic knowledge before the onset of the disease.

### Cell assemblies and word recognition procedure

Previous results obtained with analogous architectures (14, 63, 64) showed that the learning procedure described above leads to the formation of distributed associative circuits or ‘cell assemblies’ (CA, Fig.5). CAs are sets of cells that are “…more strongly connected to each other than to other” cells (65). Once developed, they behave as discrete functional units with two quasi-stable states: ‘on’ and ‘off’ (63, 64, 66). Thus, CAs are stable stimulus-specific distributed memory circuits that emerge in the network as a result of learning and exhibit complete reactivation (or ‘ignition’) in response to the presentation of the stimulus, or part of it (or even spontaneously, due to noise-driven activity accumulation (67). After the training phase (see section ‘Learning words of different semantic categories’), we applied a standard procedure to identify and quantify the cells that make up the CA circuits corresponding to each word stimulus (14, 63, 68). In the present model, spontaneously formed CAs linked word forms to aspects of those words meaning acquired through (simulated) sensory-motor areas (14).

**Fig. 5.**
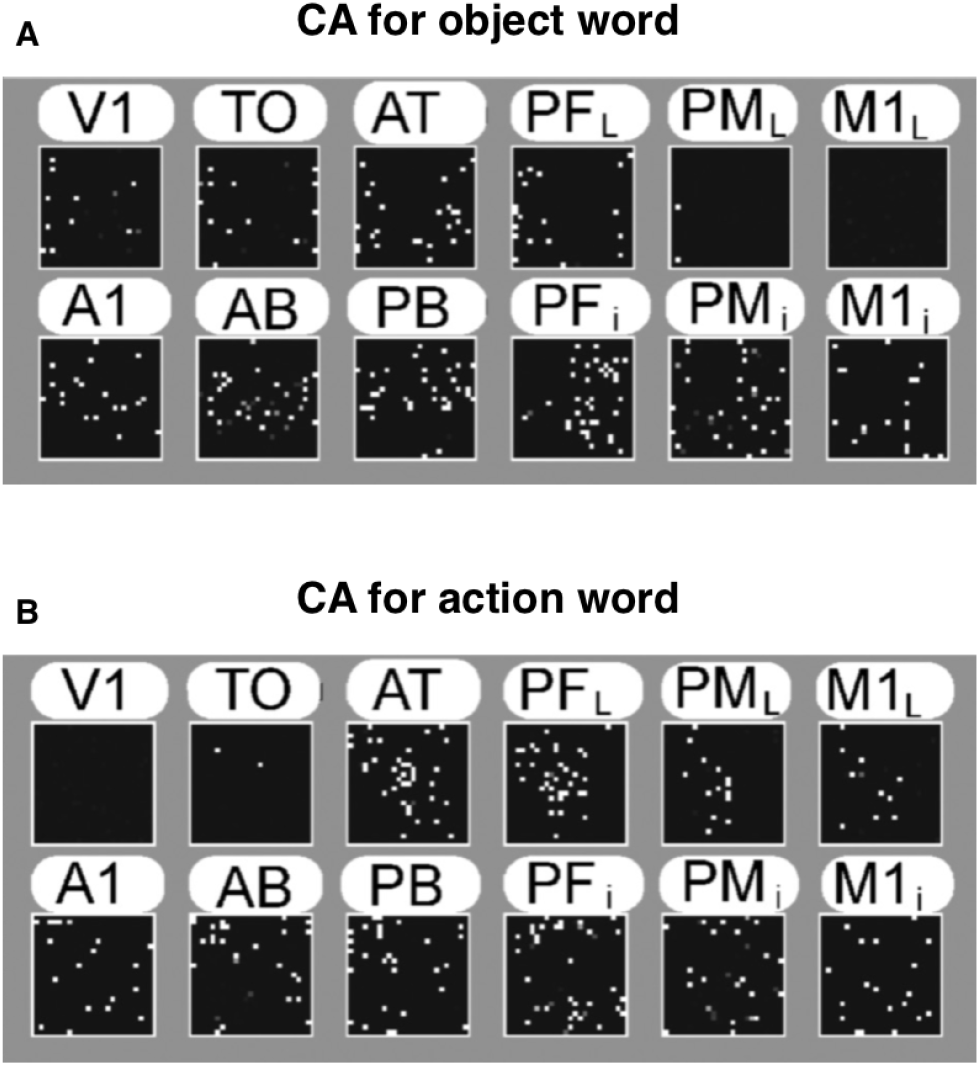
Formation of cell assembly circuits in the network as a result of simulated word learning and semantic grounding. A – example of an object-related word CA circuit. B – example of an action-related word CA circuit. Note the differential cortical distribution of the two semantic types of word circuits in the extrasylvian (but not perisylvian) areas of the model.

Crucially, before and after lesions we assessed the ability of the network to “recognize” words. To do that, the auditory component of each word pattern was presented once (for 2 time-steps) to the A1 area only while no other input was provided. The activity of each e-cell was recorded during stimulus presentation and for the following 15 time-steps. An e-cell was considered as a responsive CA cell if its time-averaged activity during this period reached a given thresh-v old (θ); this threshold is specified for each word (w) and each area (a) separately in the following way:

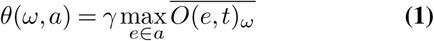

meaning that the time-averaged output is calculated for each e-cell (e) in each area (a) per each word (w). The threshold for the area for the word is equal to the fraction (γ) of the time-averaged output of the most active cell in this area for this word.

Further details of model implementation are given in the Appendix (Detailed Model Implementation).

## Results

### Main Hypothesis Testing

Figure 6 shows the number of CA cells in the extrasylvian and perisylvian areas of the network exhibiting a significant response (as defined by the recognition procedure, see ‘Methods’) as a function of SD severity. As can be seen, these results suggest that the extrasylvian portions of the CA circuits degrade more rapidly than the perisylvian portions. This difference was expected and is a direct consequence of the WM and GM lesions being applied to the AT area, which is an important component of the extrasylvian system. The qualitatively observed differential severity of degradation in the extrasylvian CA circuit was confirmed by a 2-way RM ANOVA with factors ExtraPeri (two levels — extrasylvian and perisylvian areas) and Severity (four levels — no SD, 30% SD, 60% SD, and 90% SD), revealing a significant interaction between these factors (F3,36 = 1166.29, p < 0.001).

**Fig. 6.**
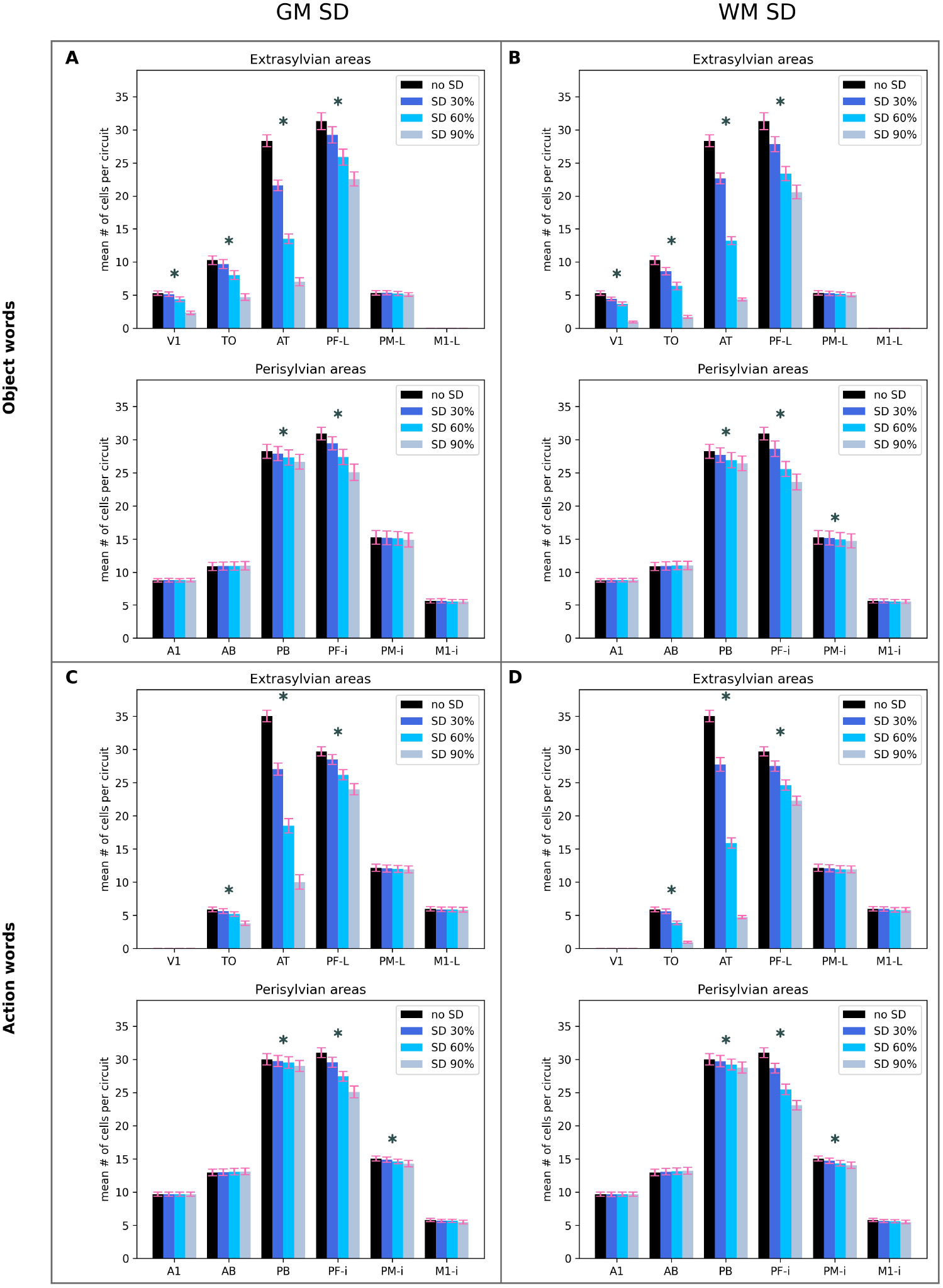
Network response during simulated auditory word recognition. The average number (in 13 simulations) of inputspecific cells exhibiting above threshold activity in response to presentation of a learned auditory stimulus (“word sound”) to the model correlate of auditory cortex is plotted as a function of area, for intact networks (no SD) and three different degrees of simulated SD lesions. A, B: responses to object words. C, D: responses to action words. A, C: simulated grey-matter (GM) SD. B, D: simulated white-matter (WM) SD. See main text for details. Error bars indicate standard error of the mean. * – difference between no SD and 90% SD cases, p<0.001, t-test.

To further explore the differences between extrasylvian and perisylvian areas, we performed an analysis of the total numbers of CA cells under those areas in four conditions: object versus action words and GM SD versus WM SD. Figure 7 shows how the number of responsive cells in the CA circuits decreases with SD progression — from no SD through 30% and 60% SD to 90% SD — in four conditions separately. Note that the data here are normalized by the number of CA cells before SD. The results show that in all four conditions the CA circuits deteriorate far more in extrasylvian areas than in perisylvian areas. For each of the four conditions, a 2-way RM ANOVA with factors ExtraPeri and Severity shows a significant interaction between factors (for four conditions: F3,36 > 307.37, p < 0.001).

**Fig. 7.**
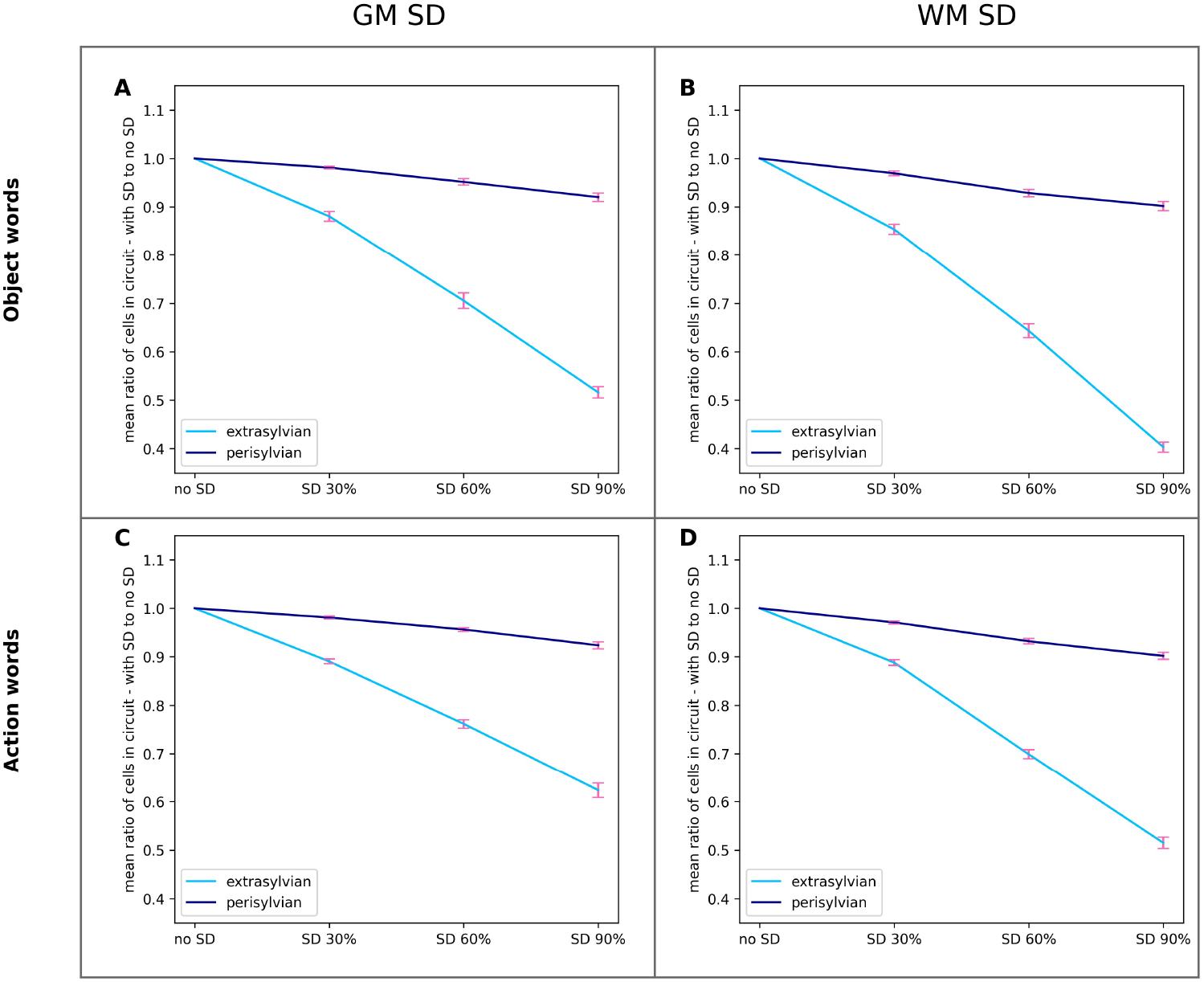
Per-system network response during simulated auditory word recognition. Same data as in Figure 6 but grouped by extra-(semantic) or peri- (phonological) - sylvian system. The ratios plotted are the percentage of word-circuit-specific reactivated cells relative to the total number of CA cells (across the system) reactivated in the intact-network (no-SD) condition. See the text for details. Error bars indicate the standard error of the mean.

Dependent samples t-tests confirm a significant difference in the number of responsive CA cells between no SD and 90% SD cases in each of the eight conditions (extra/peri x object/action x GM/WM; t12 > 10.71, p < 0.001 for all conditions). We also performed post hoc dependent samples t-tests to ensure that the number of CA cells in extrasylvian areas significantly differs from the number of CA cells in perisylvian areas in the most severe SD case (90% SD) for each of the four conditions (object/action x GM/WM; t12 > 17.88, p < 0.001). We have made a Bonferroni correction for all 12 comparisons (p-value threshold is 0.0042).

Combining this evidence, we infer that in all four conditions (object/action × GM/WM) the total number of CA cells in extrasylvian areas greatly decreases with the increase of SD severity – the average drop is 48.4% for the 90% SD case – while in perisylvian areas this number also declines, but much more slowly: the average drop is 8.7%.

To explore the SD impact on different areas, we performed post hoc dependent samples t-tests on the number of CA cells in no SD and 90% SD cases for each of the four conditions separately: object versus action and GM SD versus WM SD. All comparisons were Bonferroni corrected. Differences are presented as asterisks in Figure 6. In extrasylvian areas, SD leads to a strong decline in V1, TO, AT, and PFL areas, while PML and M1L remain intact. In perisylvian areas, SD leads to a strong decline in PFi and a mild but statistically significant decline in PB and PMi, while other areas are intact.

### Additional Findings: model predictions

We asked what predictions our model would make about recognition of object versus action words when we implemented the structural alterations reflecting the pathology of SD. (Note that we assumed here that the number of responsive CA cells can be taken as an indicator of the network’s ability to recognize words). Results shown in Fig.8 suggest that the network’s ability to respond to the presentation of the auditory part of a learned “word” declines more dramatically for object-than action-related words. The statistical validity of this observation was confirmed by the result of 2-way RM ANOVAs, ran on the total number of CA cells in extrasylvian areas in GM and WM SD, separately with factors WordType and Severity. The ANOVA for GM SD showed a significant interaction of the two factors (F3,36 = 7.04, p = 0.0012), but only a trend toward significance (F3,36 = 3.04, p = 0.067) for WM SD. However, analysis on WM SD data revealed the presence of the main effects of both WordType (F1,12 = 18.28, p = 0.001) and Severity (F3,36 = 1578.12, p < 0.001). Therefore, both for GM and WM SD, the network’s response to presentation of a known (simulated) word sound – i.e., its ability to recognize that item – decreased significantly more for object-than action-related words (48.3% decline for object words, 37.4% decline for action words, t12 = 4.61, p = 0.001 for GM SD; 59.5% decline for object words, 48.5% decline for action words, t12 = 7.82, p < 0.001 for WM SD).

**Fig. 8.**
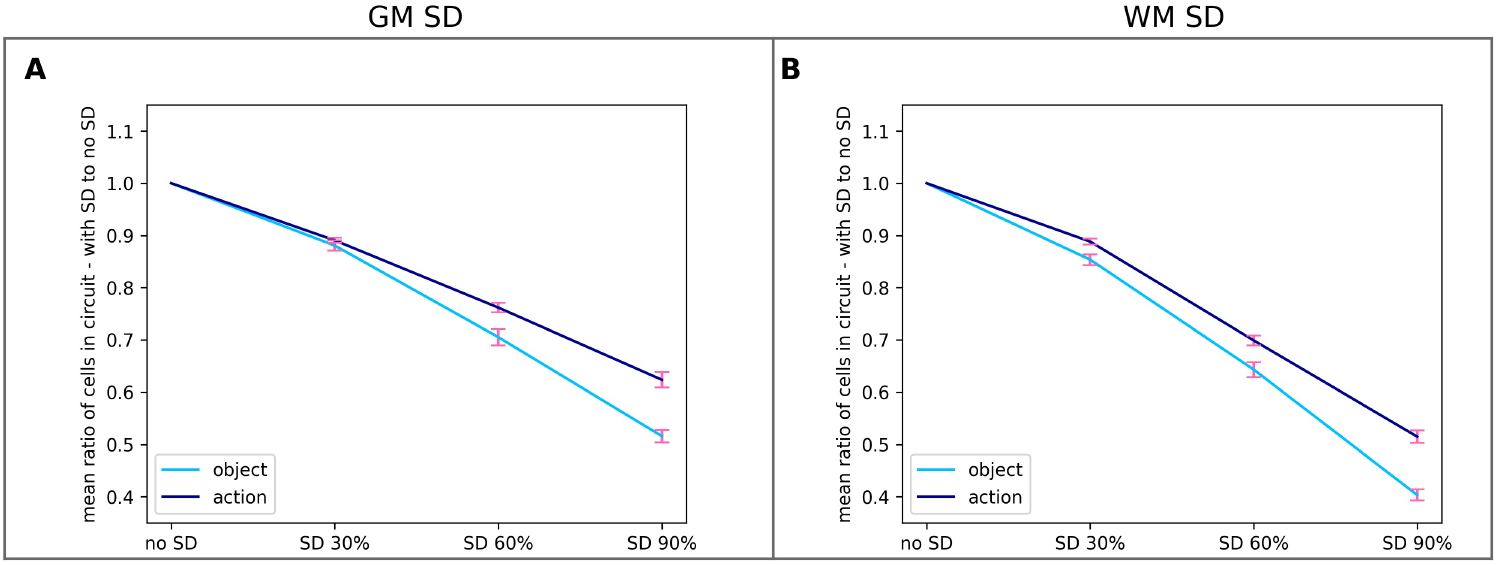
Total network response during simulated auditory word recognition for different types of words. The ratios plotted are the percentage of (word-circuit)-specific reactivated cells relative to the total number of CA cells reactivated in the intact-network (no-SD) condition. See the text for details.

We also noticed that recognition abilities appeared to decline more dramatically in WM SD than in GM SD (see Fig.9). To test this observation statistically, we ran 2-way RM ANOVAs for the total number of CA cells in extrasylvian areas (separately for object-and action-related words) with factors SD-Type and Severity. In both cases, the interaction between factors was significant (F3,36 > 28.52, p < 0.001), confirming that the network’s decrease in recognition abilities for object-and action-related words is significantly greater for WM SD than for GM SD (for object words it is 59.5% decline in WM SD vs 48.3% decline in GM SD, t12 = 8.49, p < 0.001; for action words it is 48.5% decline in WM SD vs. 37.4% decline in GM SD, t12 = 6.64, p < 0.001).

**Fig. 9.**
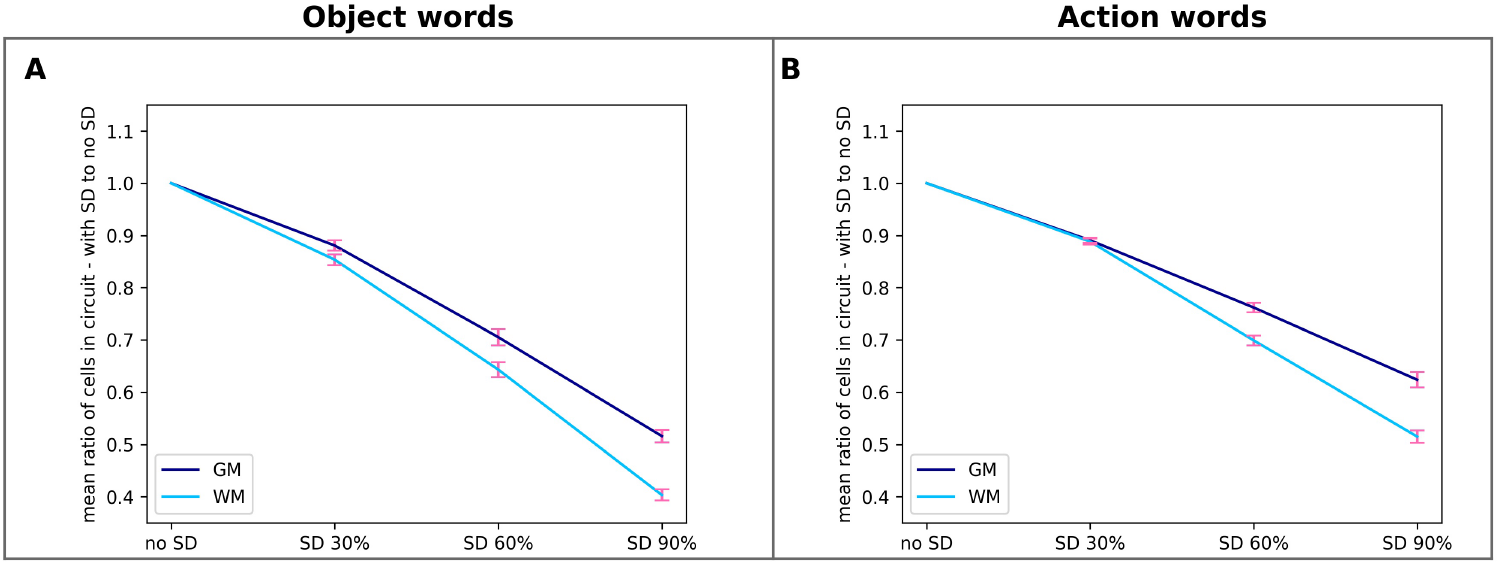
Total network response during simulated auditory word recognition for different types of SD. The ratios plotted are the percentage of (word-circuit)-specific reactivated cells relative to the total number of CA cells reactivated in the intact-network (no-SD) condition. See the text for details.

## Discussion

We used a neuroanatomically and biologically constrained neural model of the areas of the brain involved in language acquisition and semantic grounding to simulate semantic dementia (SD) lesions as progressive damage to the anterior temporal lobe – ATL –area. As a result of SD lesions, the “extrasylvian” parts of the word circuits (which include the ATL) degraded more rapidly than their perisylvian counterparts. A second striking result is the observed category-specific effects of SD lesions – namely, our simulations indicate that word processing abilities should be more impaired for object-than for action-related words; furthermore, the model shows that white matter degradations should lead to stronger deficits than grey matter lesions of analogous severity.

These simulation results are consistent with, and explain existing data about the decline of language function in clinical populations during onset and progress of semantic dementia. Specifically, the observed dissociation between word recognition deficits and relatively preserved word repetition abilities observed in SD patients is explained here by the cortically distributed character of word circuits, consisting of cell assemblies (CA) that include neurons from both extrasylvian (semantic) and perisylvian (language) areas. Previous modeling results (14) suggested that while extrasylvian areas may encode semantic knowledge and information pertaining to aspects of word meaning (and hence provide a neural substrate for word and concept recognition processes), perisylvian areas mediate the formation and storage of associations between acoustic and articulatory features of syllables and words (necessary, e.g., during word – and non-word – repetition tasks). The changes in activity patterns that we observed after introducing SD to our model speak in favor of our main hypothesis: while the CA circuits that are needed for successful word repetition (located in the perisylvian areas) remain relatively intact, word recognition abilities are progressively impaired as SD lesions increase due to a strong reduction in responsive CA cells in the extrasylvian areas. We believe that these simulated cortical changes can explain the dynamics of SD patients, who experience a strong decline of word recognition abilities but largely retain word repetition skills (10, 69, 70). We suggest that such a dissociation between perisylvian and extrasylvian areas can be further tested in neuroimaging experiments in SD patients.

We should note that the above interpretation of the model results is based on a key assumption: we are postulating that a decrease in the number of responsive extrasylvian CA cells corresponds to a de facto deficit in the word recognition ability of the network. A large body of experimental data suggests that (extrasylvian) sensorimotor areas convey information on aspects of word meaning (71–73); thus, even if parts of the CA circuits in extrasylvian areas are still activating in response to a presentation of an auditory stimulus, we submit that this partial activation contains less semantic information about the word, and therefore this information deficit translates directly into a corresponding observable word recognition deficit.

The results of the simulations also enable us to make predictions and adjudicate between competing theories on the question of whether SD patients have category-specific or category-general recognition problems. Several studies have suggested that SD leads to category-general deficits (9, 21, 74); however, our model shows that recognition of object-related words should decrease more sharply than recognition of action-related words as SD progresses. Some studies confirm that object naming is more affected in SD patients than action naming (75), and others show similar results for lesions in the ATL (76). Our model predicts (and explains) the emergence of this effect on the basis of the neuroanatomical links existing between brain areas relevant for object-and action-word processing, simulated here (see Fig.1), and specifically of the locus of the ATL itself. The ATL acts as the interface via which object-related information coming from the visual system links up with language circuits in perisylvian areas. If the grey matter within – or the white matter bundles that link to – this connector hub is lesioned, the word CA circuits that extend strongly into the visual system (i.e., object-related word circuits – see Fig.6) are severely affected, whereas CA circuits linking perisylvian and motor areas through the prefrontal hub area (PFL in the model see Fig.1), i.e., action-related word circuits, will be only marginally affected. Unfortunately, most previous studies investigating this matter tend to group words based on grammatical categories – namely, nouns or verbs – rather than on the basis of semantics properties (77); hence, to validate the above predictions, further experimental testing is desirable.

To the best of our knowledge, this is the first brain-constrained model of SD where both types of degradation white and grey matter – were implemented; previous computational studies only modeled degradation of links (WM) (15, 18). Our simulation results show that white matter degradation should lead to a more severe recognition decline than the same loss of grey matter. This additional prediction can be tested experimentally by investigating possible correlations between the grey matter / white matter loss ratio in SD and the disease severity. The results we have obtained on the distinction between grey and white matter degradation address an important feature of the cell assembly concept in our model. Although the constituent parts of CAs are individual cells, the emergence and integrity of those CAs are based on the links between them. Disabling a portion of cells naturally leads to the inactivation of some part of CA, but the same fraction of connections is more sparsely distributed and affects more cells. We can see from the results that even if not all links of some particular cell are inactivated – which would be the same as disabling the cell itself – those that are could be crucial for the cell’s functioning as a part of CA.

Another aim of this paper has been to argue computationally that hybrid theories of semantic system organization offer the most promising theoretical approach to explain a wide range of experimental data. The present model can be considered an example of such hybrid paradigms: the primary areas in the action and perception brain systems are particularly important to provide sensorimotor co-experiences during word meaning acquisition; the central (semantic hub) areas enable these experiences to converge and be associated together into a single (word, or conceptual) circuit; the emergent semantic representations are distributed between the modality-specific sensorimotor areas and convergence hubs. One of the arguments of proponents of amodal theories is that ATL must be the place where concepts are stored in an amodal way because its degradation leads to semantic system deterioration (9, 21). However, as we show here, if word circuits are distributed across multiple diverse areas (all of which together support word recognition and processing), ATL damage can still lead to semantic impairments as observed in SD without this hub necessarily being the locus of amodal conceptual representations.

To conclude, the present neurobiologically constrained model explains – in terms of cortical mechanisms and neuroanatomical characteristics – the existing data on language impairments as observed in SD clinical populations. Furthermore, for the first time, simulated white or grey matter lesions to the anterior-temporal lobe area of the model enable making novel, theory-driven predictions about (i) differential effects of semantic dementia on word processing deficits (specifically, on object-vs action-related word recognition) and (ii) different degrees of language impairment depending on the type of tissue damaged by the neurodegenerative disease.

## Supplementary Note 1: Appendix (Detailed Model Implementation)

### Microstructure

Each area consists of two neuronal layers — an excitatory layer and an inhibitory one – with 625 (25×25) cells (e-cells for the former and i-cells for the latter) in each layer. To avoid any potential edge effects, layers have a toroidal structure: the top edge is adjacent to the bottom one, and the left edge is adjacent to the right one. I-cells and e-cells have one-to-one correspondence. A combination of an e-cell and an i-cell reflects a population of pyramidal excitatory neurons and inhibitory interneurons of one cortical column (grey matter under approximately 0.25 mm2 of the cortical surface). Cells are modeled as graded-response neurons.

Each e-cell is restricted to send its projections to the 19×19 e-cell patch in the same area, 19×19 e-cell patches in connected areas, and 5×5 i-cell patch in the same area (Fig.2). These projections are created during the first initialization of a network in a random manner. The probability that the projection is created is derived from the Gaussian probability density function (Table 2). The highest probability is in the patch center (although, a cell cannot send projection to itself) and it is decreasing with the distance from the center.

**Table 1.**
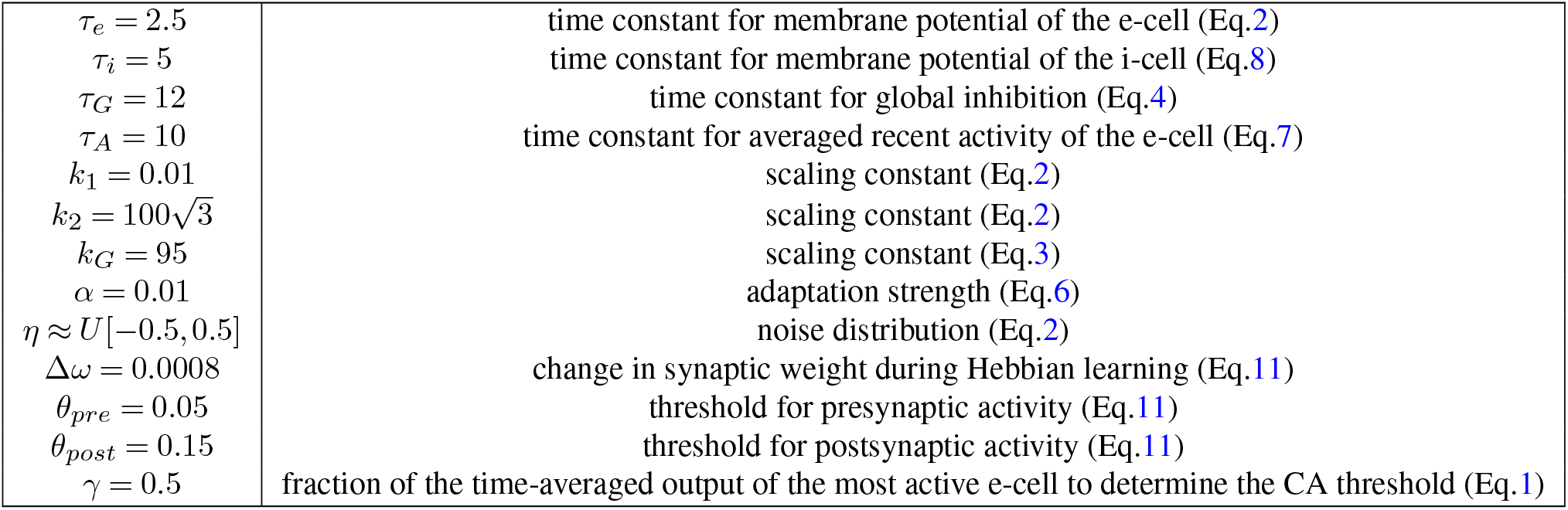
Parameters of the model.

**Table 2.** Gaussian distribution parameters. The probability that links between two cells will be created follows the Gaussian distribution, decreasing with the distance between cells.

|  |  |
| --- | --- |
| $P_{rec} = 0.15$ | central probability of the Gaussian distribution for recurrent links |
| $P_{between} = 0.28$ | central probability of the Gaussian distribution for links between different areas |
| $\sigma_{rec} = 4.5$ | standard deviation for the Gaussian distribution for recurrent links |
| $\sigma_{between} = 6.5$ | standard deviation for the Gaussian distribution for links between different areas |
| $\omega_{initial} \approx U[0, 0.1]$ | initial weight distribution for links between two e-cells |
| $\omega_{inh} \approx N[0.295, 0.2]$ | weight distribution for links from the e-cell to a 5x5 patch of i-cells, decreasing with the distance between cells |

### E-cell dynamics

The activity state of a cell is uniquely defined by its membrane potential. The change in the e-cell membrane potential is dependent on its current membrane potential, a sum of its inputs, and uniform distribution of white noise:

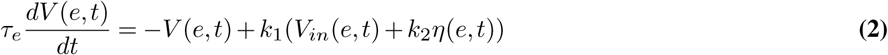

where *V* (*e, t*) is a membrane potential of the E-cell

*V*_*in*_(*e, t*) is a sum of inputs to this E-cell (see equation 3)

*η*(*e, t*) is noise

*τ*_*e*_ is the time constant

*k*1 and *k*2 are the scaling constants

(Membrane potential of the e-cell)

Note that each e-cell has noise as its property which represents the spontaneous activity of real neurons. The sum of inputs to the e-cell is governed by the equation:

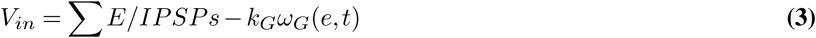

where ∑ *E/IPSPs* is the sum of excitatory and inhibitory postsynaptic potentials sent to the cell of interest

*ω*_*G*_(*e, t*) is global inhibition (see equation 4)

*k*_*G*_ is the scaling constant

(Sum of inputs to the e-cell)

Postsynaptic potential is defined as the output of the cell (see equations 5 and 10) multiplied by the synaptic weight between this cell and the target cell. Each e-cell gets exactly one IPSP from the corresponding i-cell, which is included in this sum with a negative sign. Note the global inhibition mechanism that is an area-specific inhibitory loop that prevents overall network activity from falling into nonphysiological states. This inhibition is calculated as follows:

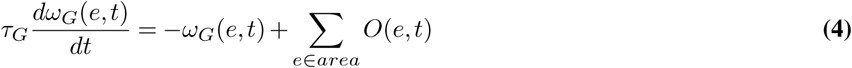

Where∑_*e ∈area*_ *O*(*e, t*) is the sum of e-cells’ outputs from the whole area (see equation 4)

*τ*_*G*_ is the time constant

(Global inhibition of the E-cell)

An output of the e-cell is some value between zero and one. It depends on the cell’s membrane potential and the threshold that this potential needs to exceed for an output to have a non-zero value:

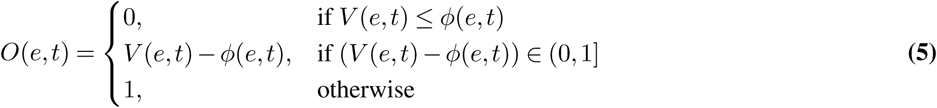

where *O*(*e, t*) is the output of the e-cell

*ϕ*(*e, t*) is the threshold of the membrane potential (see equation 6)

(Output of the e-cell)

The threshold of the membrane potential is not constant, it depends on the cell’s recent activity – the more active the cell was, the higher is the threshold:

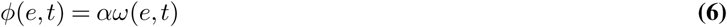

where *ω*(*e, t*) is averaged recent activity of an e-cell (see equation 7)

*α* is adaptation strength

(Threshold of the membrane potential)

When the network is initiated, the recent activity of every cell is zero (*ω*(*e, t*) = 0). Then it is updated on each time step with an increase/decrease calculated as follows:

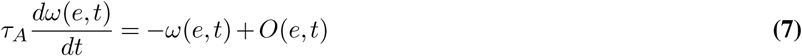

where *τ*_*A*_ is the time constant

(Averaged recent activity of the e-cell)

### I-cell dynamics

Change in the i-cell membrane potential is calculated similarly to those of the E-cell, but without the white noise component:

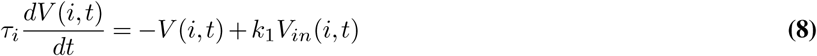

where *V* (*i, t*) is the membrane potential of the I-cell

*V*_*i*_*n*(*i, t*) is the sum of inputs to this I-cell (see equation 9)

*τ*_*i*_ is the time constant

*k*_1_ is the scaling constant

(Membrane potential of the i-cell)

There is no global inhibition mechanism for the i-cell as there is for the e-cell and its sum of inputs is just the sum of EPSPs it received:

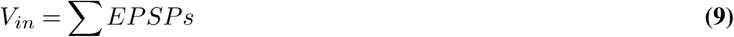

where ∑ *EPSPs* is the sum of EPSPs to the i-cell (Sum of the inputs to the i-cell)

Note that the i-cell can receive only excitatory inputs from the 5×5 e-cells patch.

The output of the i-cell is simply its membrane potential if it is positive or zero otherwise:

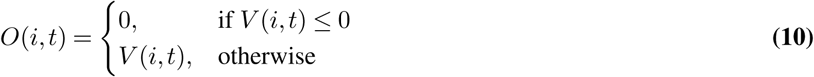

where *O*(*i, t*) is the output of the i-cell

### Synaptic weight dynamics

Initially, random weights are assigned to all established connections with the weights of the links between two e-cells distributed uniformly and the weights of the links from e-cells to i-cells distributed normally, decreasing with the distance between the cells. Then, those weights are dynamically changing via the Hebbian learning mechanism:

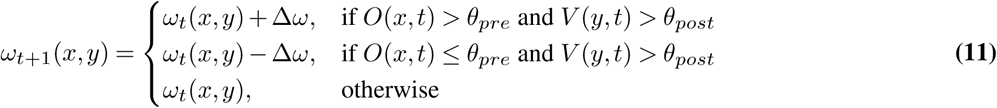

where *ω*_*t*_(*x, y*) is the weight of the synapse from the cell *x* to the cell *y* at time *t*

Δ*ω* is the change in the synaptic weight

*θ*_*pre*_ is the presynaptic output activity threshold

*θ*_*post*_ is the postsynaptic membrane potential threshold

If both the output of the presynaptic cell and the membrane potential of the postsynaptic cell are high enough, synaptic weight is increased by constant number (long-term potentiation, LTP), which corresponds to the rule “fire together — wire together”. If low presynaptic output is correlated with high postsynaptic potential, synaptic weight is weakened by the same amount (long-term depression, LTD), which corresponds to the rule “out of sync— out of link”.

## Notes

https://github.com/ansty57/Master_Thesis_hse2020

### Competing Interest Statement

The authors have declared no competing interest.

### Summary of Updates

Authors list updated

